# SARS-CoV-2 spike glycosylation affects function and neutralization sensitivity

**DOI:** 10.1101/2023.06.30.547241

**Authors:** Fengwen Zhang, Fabian Schmidt, Frauke Muecksch, Zijun Wang, Anna Gazumyan, Michel C. Nussenzweig, Christian Gaebler, Marina Caskey, Theodora Hatziioannou, Paul D. Bieniasz

## Abstract

The glycosylation of viral envelope proteins can play important roles in virus biology and immune evasion. The spike (S) glycoprotein of severe acute respiratory syndrome coronavirus-2 (SARS-CoV-2) includes 22 N-linked glycosylation sequons and 17 O-linked glycosites. Here, we investigated the effect of individual glycosylation sites on SARS-CoV-2 S function in pseudotyped virus infection assays and on sensitivity to monoclonal and polyclonal neutralizing antibodies. In most cases, removal of individual glycosylation sites decreased the infectiousness of the pseudotyped virus. For glycosylation mutants in the N-terminal domain (NTD) and the receptor binding domain (RBD), reduction in pseudotype infectivity was predicted by a commensurate reduction in the level of virion-incorporated spike protein. Notably, the presence of a glycan at position N343 within the RBD had diverse effects on neutralization by RBD-specific monoclonal antibodies (mAbs) cloned from convalescent individuals. The N343 glycan reduced overall sensitivity to polyclonal antibodies in plasma from COVID-19 convalescent individuals, suggesting a role for SARS-CoV-2 spike glycosylation in immune evasion. However, vaccination of convalescent individuals produced neutralizing activity that was resilient to the inhibitory effect of the N343 glycan.

## Introduction

Severe acute respiratory syndrome coronavirus 2 (SARS-CoV-2) is the causative agent of the COVID-19 disease and has caused a devastating pandemic (1, 2). SARS-CoV-2 encodes a spike (S) glycoprotein which binds angiotensin-converting enzyme 2 (ACE2) and mediates viral entry into host cells (3-6). The S glycoprotein (1273 aa) consists of a signal peptide followed by the S1 subunit (13-685 aa) and the S2 subunit (686-1273 aa). These two subunits are separated by a furin cleavage site (PRRAR), abrogation of which can increase virus infectivity in some circumstances (7). The receptor-binding domain (RBD, 319-541 aa) that is responsible for ACE2 binding (6) and the N-terminal domain (NTD), both encoded in S1, are the major targets of neutralizing antibodies. Like other viral envelope glycoproteins including HIV-1 (8), SARS-CoV-2 spike protein is heavily glycosylated (9-11). Indeed, approximately 40% of the surface the SARS-CoV-2 S protein expressed in human 293T cells is shielded by glycans (12). The majority of this shield is comprised of N-linked oligomannose-type or complex glycans, linked to 22 sites (Asn-X-Ser/Thr) on the S protein (13). Additionally,17 O-linked glyco-sites have been identified by biochemical methods (14-16).

Glycosylation of viral envelope proteins can play an important role in virus-host interactions (17). In the case of HIV-1, for example, N-linked glycans are essential for correct folding and processing of gp120 as well as structural rearrangements required for receptor binding (18). The HIV-1 glycan shield also plays a crucial role in preventing neutralizing antibodies from binding to HIV-1 envelope (19). Likewise, S glycosylation affects SARS-CoV-2 infection (20). Blocking N-glycan biosynthesis onto SARS-CoV-2 spike protein and, to lesser extent, O-glycan elaboration, reduces viral infectivity (21). Additionally, cryo-electron microscopy studies have revealed that the N-glycan at position N343 in the RBD facilitates transition of the spike protein to the ‘open’ conformation, which is important for ACE2 binding (22). Accordingly, mutation of this site (N343Q) reduced viral entry into ACE2-expressing cells (23).

Little is known about how SARS-CoV-2 S glycosylation might affect immune surveillance. It is conceivable that glycans sterically shield underlying epitopes from recognition by antibodies, as is the case in HIV-1 (19). Many SARS-CoV-2 neutralizing antibodies target the RBD and can be divided into 4 broad classes based on the epitopes targeted (24). Class 1 antibodies recognize epitopes overlapping with the ACE2-binding site and bind only to ‘up’ RBDs. Class 2 antibodies bind both ‘up’ and ‘down’ RBDs and also block ACE2 binding. In contrast, class 3 antibodies bind epitopes distinct from the ACE2 binding site but can potently neutralize. Class 4 antibodies are generally less potent and recognize epitopes that are distinct from the ACE2 binding site that are shielded in the down conformation. Interestingly, some antibodies, namely S309 and SW186, recognize epitopes that include the N343 glycan (25) (26), raising the possibility that this glycan might play a dual role in antibody recognition, either as shield or as a component of an epitope.

Co-expression of SARS-CoV-2 S with envelope-deficient viruses such as HIV-1 (human immunodeficiency virus-1) produces pseudotyped viruses capable of infecting ACE2-expressing cells and is widely used as a surrogate to study viral entry and neutralization by antibodies (27, 28). To comprehensively understand the role of spike glycans in viral infectivity and antigenicity, we individually mutated each of the 22 N-linked glycosylation sites in the spike protein as well as two O-linked sites (S323 and T325) in the RBD. We found mutations introduced at many glycosylation sites in the NTD and RBD reduced pseudotype infectivity, and the magnitude of this effect was predicted by the magnitude of the loss of S incorporation into virions. Furthermore, while the S protein levels in virions had little effect on neutralization sensitivity, the presence or absence of a glycan on N343 in RBD governed the sensitivity to some monoclonal antibodies cloned from convalescent individuals. Moreover, the glycan at N343 reduced neutralization sensitivity to polyclonal antibodies from convalescent individuals, but this evasive effect imparted by glycosylation was overcome by plasma antibodies from the same individuals who were subsequently vaccinated.

## Results

### Levels of virion-incorporated SARS-CoV-2 spike protein and pseudotyped HIV-1 particle infectiousness

While SARS-CoV-2 S pseudotyped HIV-1 has been widely utilized to study S-mediated viral entry and neutralizing activities by antibodies, the extent to which the amount of S protein on virions affects particle infectivity and neutralization sensitivity is not fully understood. To address this question, we co-transfected 293T cells with various amounts of an S expression plasmid (pSARS-CoV-2_Δ19_), along with an envelope-deficient HIV-1 proviral plasmid encoding NanoLuc. The number of SARS-CoV-2 spikes incorporated into HIV-1 pseudotype virions were estimated using purified recombinant purified S protein (S-6P-NanoLuc (29)) and recombinant p24 CA proteins as standards on near-IR fluorescent western blotting (LiCor) and assuming 1500 capsid (CA) protein subunits per mature HIV-1 particle (30, 31) (Fig. S1A). While the amount of S expression plasmid used for transfection had little effect on the amount of virions produced (Fig. 1A), the amount of S protein incorporated into virions correlated with the amount expressed in cells. S incorporation into virions reached a plateau (∼30 ng S/ml in virions, or around 300 spike trimers per virion) when 0.125 μg or 0.25 μg of an S expression plasmid was co-transfected with the HIV-1 proviral plasmid (Fig. 1A, 1B, 1C).

**FIG 1.**
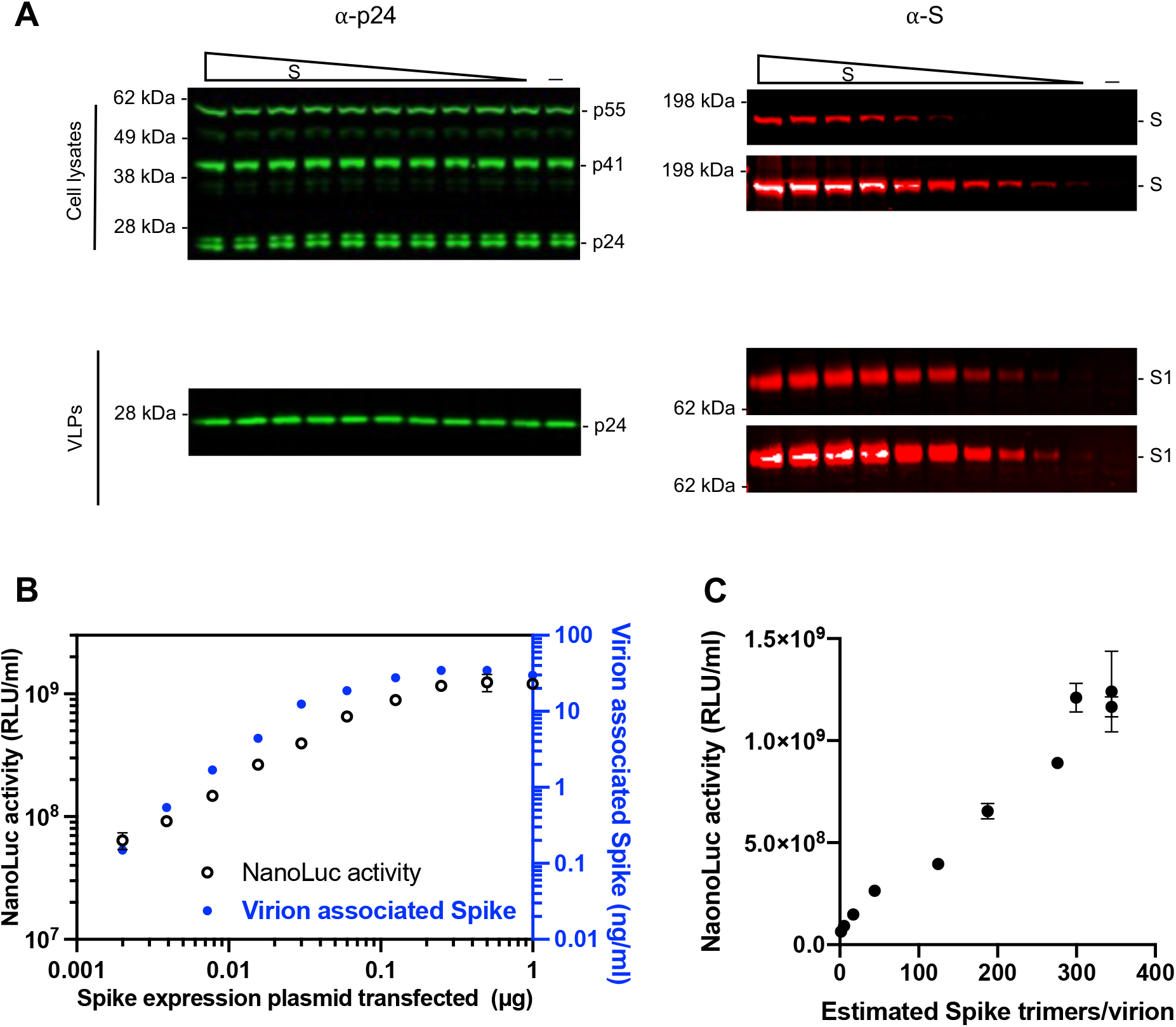
The effect of spike proteins level on pseudotype infectiousness. (A) Western blot analysis of 293T cell lysates (upper panel) or virions (lower panel) at 48 hours after transfection with various amounts (0 ng, 2 ng, 3.9 ng, 7.8 ng, 15.6 ng, 31.2 ng, 62.5 ng, 125 ng, 250 ng, 500 ng, or 1000 ng) of a WT SARS-CoV-2 spike expression plasmid along with envelope-deficient HIV-1 proviral plasmid expressing NanoLuc luciferase. Each S blot was scanned twice, at low intensity (*upper*) and high intensity (*low*er), respectively. Representative of three independent experiments. (B) Characterization of virions pseudotyped with spike expression plasmids. Infectiousness (on left y axis) was quantified by measuring NanoLuc luciferase activity (RLUs) following infection of 293T cells expressing ACE2 (293T/ACE2.cl22) in 96-well plates with pseudotyped viruses. The S1 incorporated into viruses (on right y axis) was determined by quantitative Western Blot, using purified recombinant S-6P-NanoLuc proteins as standard, representative of three independent experiments. The mean and range of two technical replicates are shown. (C) Characterization of virions pseudotyped with spike expression plasmids as in (B). virus infectiousness (on y axis) was plotted against S trimers per virion (on x axis), which was determined by quantitative Western blot, using purified recombinant S-6P-NanoLuc proteins as standard. Representative of three independent experiments.

To assess the effect of the number of spikes on pseudotype infectiousness, the titers of pseudotyped viruses were measured on 293T cells expressing ACE2 (293T/cl.22). Co-transfection with as little as 2 ng of S expression plasmid produced virions that induced 1000x the level of luciferase observed with ‘bald virions’, indicating that small amounts of S protein (0.15 ng/ml or a mean of ∼1-2 S trimers per virion) were sufficient to mediate viral entry (Fig. S2, Fig. 1B and 1C). Nevertheless, pseudotype virion infectivity increased with increasing spike numbers. Indeed, infectivity and the number of spikes was approximately linearly correlated, in the range 1 spike to 300 spikes per virion (Fig. 1C). Of note, these spike numbers are comparable to the average number of S trimers on authentic SARS-CoV-2 virions (25–127 prefusion spikes per virion) (32) and higher than the numbers of gp120 trimers on HIV-1 virions (33).

### Reduced infectious virion yield conferred by SARS-CoV-2 spike glycosylation site removal

To investigate the contribution of glycosylation to S protein function, we generated 22 spike substitution mutants, each containing a single Asn to Asp (N to D) substitution at one of the 22 N-glycosylation sequons. Additionally, we generated a mutant with Ala replacements at potential O-linked sites S323 and T325 in the receptor binding domain (RBD) (13). None of these substitutions affected the levels of the S protein in transfected cells, but some of them reduced S incorporation into virions (Fig. 2A). Specifically, N61D, N122D, N165D, N343D, and S323A T325A that fall within S1, that includes the NTD and RBD, exhibited 10-fold or greater reductions in the levels of virion-associated S protein, suggesting that these glycans affect S protein transport or virion incorporation. Conversely, removal of the glycosylation sites in S2 had either no or minor effects on S protein incorporation into virions (Fig. 2A). Measurements of the yield of infectious pseudotyped particles carrying each of the substitutions indicated that several substitutions in the NTD and RBD markedly reduced infectivity. For example, the N61D substitution reduced infectivity by almost 50-fold (Fig. 2B, Fig. S2) while substitutions at glycosylation sites in the RBD, namely N331D, N343D, and S323A T325A, resulted in 5- to 10-fold reduction in infectivity (Fig. 2B, Fig. S2). Western blot analyses revealed that the amount of S protein in virions was directly correlated with infectivity (Fig. 2B), suggesting a potential role for most glycans during synthesis or folding of spike trimers, or their incorporation into virions, rather than in S protein function after virion incorpration. In contrast, two substitutions (N1194D and N657D) did not change the amount of S protein in virions, but substantially reduced infectivity (Fig. 2B, Fig. S2). This finding suggests a functional role for N1194 and N657 glycans in spike conformation or stability on virions, or function during viral entry. Overall, we conclude that a subset of glycosylation sites is important for the incorporation of S protein into virions, while others are required for optimal particle infectivity.

**FIG 2.**
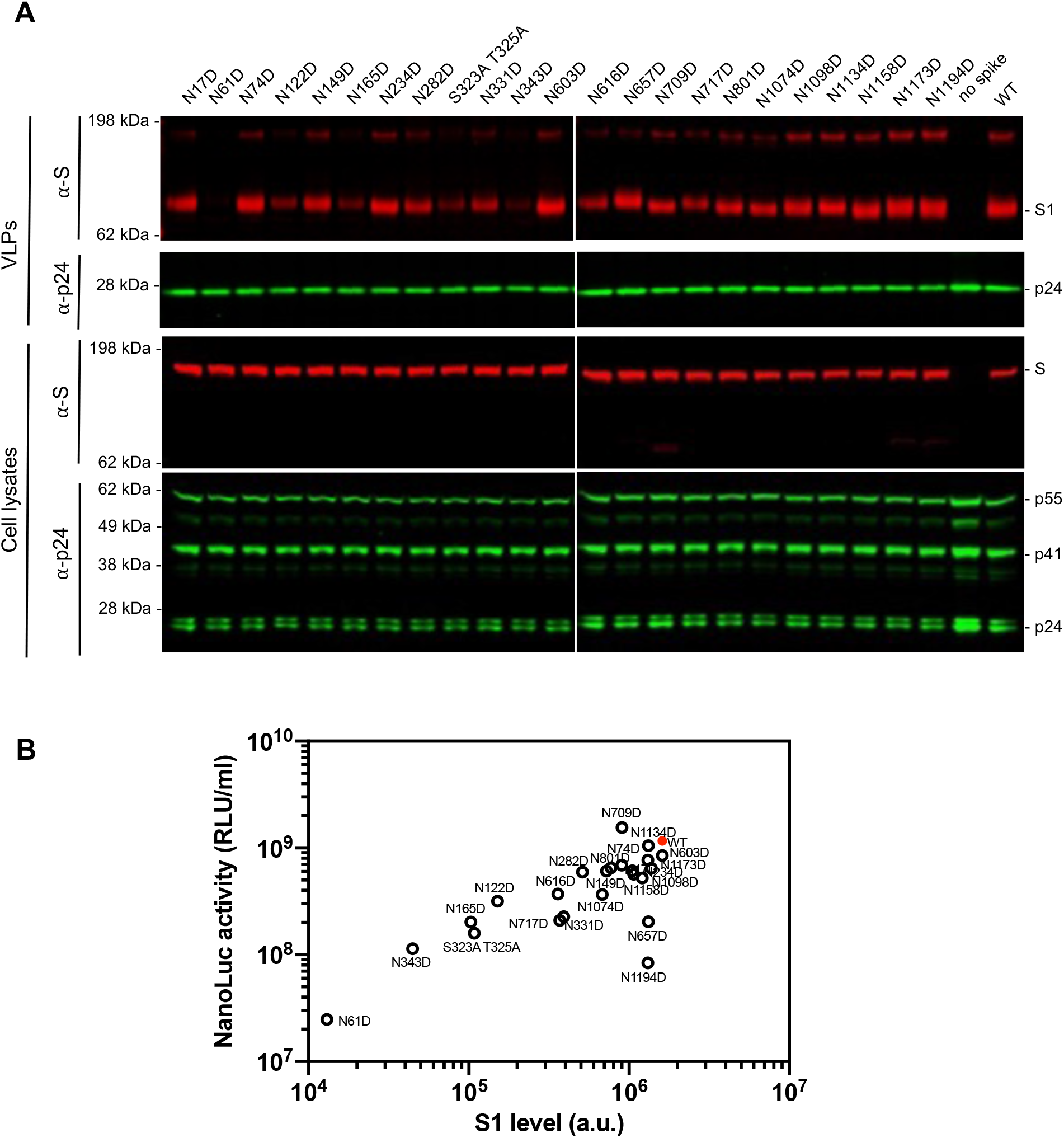
The impact of glycosylation site mutations on spike incorporation and particle infectivity. (A) Western blot analysis of 293T cell lysates (lower panel) or virions (upper panel) at 48 hours after transfection with 1 μg of glycosylation site mutants or wild-type spike expression plasmid along with envelope-deficient HIV-1 proviral plasmid expressing NanoLuc. Representative of two independent experiments. (B) Characterization of virions pseudotyped with spike expression plasmids. Infectiousness (on y axis) was quantified by measuring NanoLuc luciferase activity (RLUs) following infection of 293T/ACE2.cl22 cells in 96-well plates with pseudotyped viruses. The S1 incorporated into viruses (on x axis) was determined by quantitative Western blot. Glycosylation site mutants are shown in black and wild-type spike is shown in red. The mean and range of two technical replicates are plotted.

### Impact of spike density and RBD glycosylation on neutralization sensitivity

he RBD is the major target of neutralizing antibodies. To determine the effect of glycan removal on sensitivity to neutralizing antibodies, we focused on glycosylation sites in the RBD (N331 and N343), and one site adjacent to the RBD (N282). Because these glycosylation sites affected the level of spike incorporation into virions, we first tested the effect of SARS-CoV-2 spike density on neutralization sensitivity, as this parameter could be a potential confounder. We harvested pseudotyped virions from cells transfected with varying amounts (from 2 ng to 1 μg) of wild-type S expression plasmid and tested their sensitivity to C144, a potent class 2 neutralizing human monoclonal antibody cloned from a convalescent individual (34) (Fig. 3A). Notably, varying the levels of WT S protein on virions over a wide range (Fig. 1B) had no discernable effect on neutralization sensitivity to C144.

**FIG 3.**
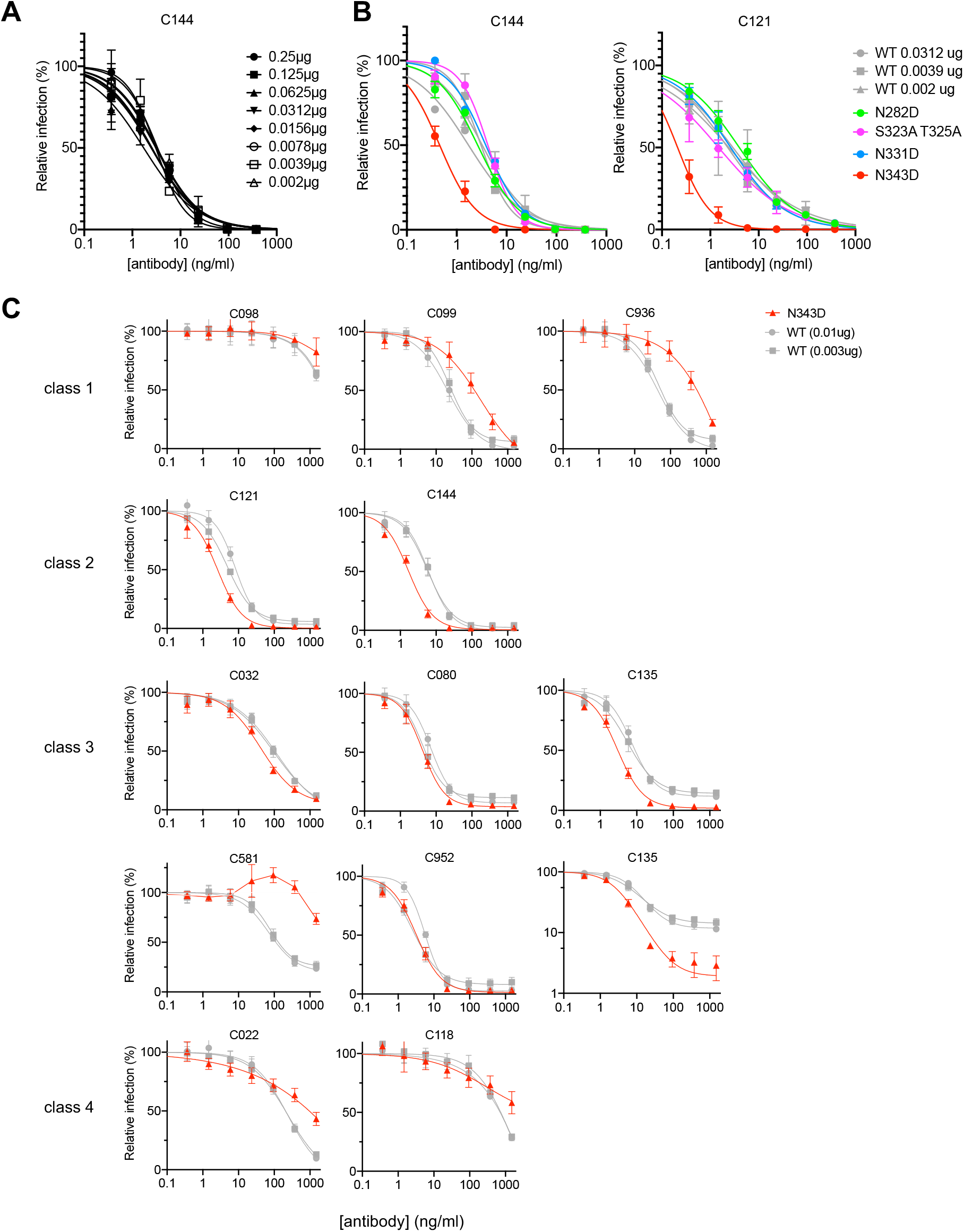
Neutralization sensitivity of Spike to monoclonal antibodies is affected by N343D substitution. (A) Quantification of pseudotyped virions from cells transfected with various amounts of wild-type spike expression plasmid along with envelope-deficient HIV proviral plasmid expressing NanoLuc in the presence of the indicated concentrations of the class 2 neutralizing monoclonal antibody C144. Infectivity was quantified by measuring NanoLuc luciferase levels (RLU). The mean and range of two technical replicates are shown. (B) Quantification of glycosylation site mutant N282D, S323A T325A, N331D, and N343D or wild-type spike pseudotyped virus infection in the presence of the indicated concentrations of the class 2 neutralizing monoclonal antibody C144 (*left*) or C121 (*right*). Infectivity was quantified by measuring NanoLuc luciferase levels (RLU). The mean and range of two technical replicates are shown. (C) Quantification of neutralization of glycosylation site mutant N343D in the background of furin uncleavable (R683G) SARS-CoV-2 S pseudotyped virus infection in the presence of the indicated concentrations of a panel of monoclonal antibodies, including class 1 (C098, C099, and C936), class 2 (C121 and C144), class 3 (C032, C080, C135, C581, and C952), and class 4 (C022 and C118) antibodies. As controls, glycosylation intact spike expression plasmid (WT in the R683G background) was transfected at two doses, 10 ng or 3 ng, and the neutralization sensitivity of the resulting viruses were assessed in parallel. The mean and range of two technical replicates are shown. The C135 neutralization graph is depieced with both a linear and logarithmic Y-axis to more clearly show effects of glycosylation on the completeness of neutralization at high antibody concentrations

We next tested the neutralization sensitivity of the RBD glycosylation site mutants bearing Asn to Asp mutation at N-linked sites (N331D and N343D) or proximal (N282D) to the RBD or alanine substitutions at O-linked sites S323 and T325. To provide matched control viruses with approximately similar numbers of S trimers and similar levels of infectivity, neutralization of the glycosylation site mutants was compared to virions generated with an appropriate level of the WT spike protein. We compared the neutralization sensitivity of WT and mutant virions to C144 and another potent class 2 neutralizing antibody, C121, both of which target the ACE2 binding site. While the N282D, S323/T325A, and N331D mutant pseudotypes exhibited neutralization sensitivities that were similar to the WT pseudotype, the N343D substitution conferred significantly increased neutralization sensitivity to both C144 and C121 (Fig. 3B). Specifically, the half maximal inhibitory concentration (IC_50_) of C144 was reduced from 1.8 to 3.4 ng/ml (against the WT pseudotype) to 0.45 ng/ml (against the N343D pseudotype), while the C121 IC_50_ was reduced from 2.3 to 2.9 ng/ml (against the WT pseudotype) to 0.21 ng/ml (against the N343D pseudotype).

### Positive and negative effects of RBD glycosylation on sensitivity to human monoclonal antibodies

To determine the effects of glycosylation more broadly on SARS-CoV-2 sensitivity to neutralizing antibodies, we used pseudotyped viruses bearing spike proteins with R683G substitution, which ablates the furin cleavage site. This substitution does not affect S incorporation into virions (Fig. S1B) but enhances particle infectivity (7). The glycosylation site mutations had a smaller effect on spike incorporation into virions in the R683G context (Fig. S3A). Nevertheless, transfection of cells with 1 μg of N282D, S323A T325A, and N331D S expression plasmids generated pseudotyped viruses with S protein amounts and infectious properties comparable to those generated by transfection of 30-100 ng of WT S expression plasmid (Fig. S3A and S3B). Transfection of cells with 1 μg of the N343D spike expression plasmid yielded pseudotyped virus particles carrying a similar amount of S protein to those generated by cells transfected with 10 ng of the R683G S protein expression plasmid (Fig. S3A), while the particle infectivity was similar to those from cells transfected with 3 ng of the WT R683G expression plasmid (Fig. S3B). To evaluate the effect of glycosylation site mutations on sensitivity to neutralizing antibodies, we generated WT and mutant virion stocks bearing similar amounts of spike protein and then assessed their susceptibility to neutralization by a panel of RBD-specific human monoclonal antibodies of the various neutralizing classes (24) recovered from convalescent individuals, including antibodies from class 1 (C098 (35), C099 (35), C936 (36)), class 2 (C121 (34), C144 (34)), class 3 (C032 (34), C080 (35), C135 (34), C581 (36), C952 (24)), and class 4 (C022 (34) (37), C118 (34) (37)).

Of class 1 antibodies, C098 had only weak neutralizing activity, whereas its clonal relative C099 was potent (IC_50_ =21.3 ng/ml against WT (10ng) or 24.5 ng/ml against WT (3ng) and its activity was resilient to many naturally occurring mutations in the RBD (7). The N343D mutation decreased the neutralization sensitivity to C099 by 7-fold, ie IC_50_ was increased to 176.9 ng/ml (Fig. 3C). Neutralization sensitivity to a third unrelated class 1 antibody, C936, was reduced by almost 10-fold, and IC_50_ was increased from 41.1 ng/ml against WT (10ng) or 54.9 ng/ml against WT (3ng) to 467.1 ng/ml against N343D (Fig. 3C).

In contrast to the class 1 antibodies, pseudotyped virus sensitivity to two class 2 antibodies was increased for the N343D mutant compared to WT. As was the case in the context of the furin-cleavable S protein (Fig. 3B), the N343D mutation in R683G spike increased neutralization sensitivity to C121 and C144. The IC_50_ of C121 was reduced from 4.8 to 8.1 ng/ml (against the WT) to 2.6 ng/ml (against the N343D), while the C144 IC_50_ was reduced from 5.7 to 6.1 ng/ml (against the WT) to 1.7 ng/ml (against the N343D) (Fig. 3C).

Class 3 antibodies, which bind epitopes distinct from the ACE2 binding site, showed a complicated pattern of effects in response to the N343D substitution. Some class 3 antibodies, including C032, C080, and C952, inhibited the WT and N343D mutant pseudotypes with approximately the same potency. A different class 3 antibody C135, that exhibited incomplete neutralization of the WT pseudotypes at high concentrations despite exhibiting low IC_50_ (IC_50_= 8.3 -11.5ng/ml), was able to achieve almost complete neutralization of the N343D pseudotypes at high concentrations (Fig. 3C). In contrast, another class 3 antibody C581 showed reduced potency against the N343D mutant, ie, the IC_50_ was >2000 ng/ml against N343D compared to 70.0 to 76.6 ng/ml against the wild-type pseudotype (Fig. 3C).

For two class 4 antibodies, C022 and C118, the N343D substitution affected the slope of the neutralization curves and conferred partial resistance at high of antibody concentrations (Fig. 3C). For example, C022 at 2000 ng/ml almost completely neutralized the wild-type pseudotypes but only inhibited the N343D pseudotypes by ∼50%. A similar trend was seen for a second class 4 antibody, C118.

Overall, the N343D substitution had a range of positive and negative effects on neutralization sensitivity that varied greatly dependent on the nature of the particular monoclonal antibody tested. To better understand the molecular basis for the impact of N343 glycan on antibody neutralization, we inspected the structures of some of the aforementioned antibodies in complex with spike (Fig. 4A, B, C). The glycosylation site at N343 (shown in red in Fig. 4A, B, C) is distinct from the binding site of the class 1 antibody C098 – suggesting that the effects of the glycan on sensitivity to class 1 antibodies are mediated through effects of the glycan on RBD conformational dynamics and epitope exposure (Fig. 4B). In contrast, for the class 2 antibodies C121 and C144, N343 is proximal to the antibody bound to the neighboring spike subunit (Fig. 4C). Since removal of the glycan increased sensitivity to these antibodies (Fig. 3C), it is likely that the N343 glycan partly shields these class 2 antibody epitopes. For class 3 antibodies, N343 protrudes towards the C135 binding site on the same S subunit (Fig. 4), potentially explaining incomplete neutralization by this antibody (Fig. 3C). For the class 4 antibody C118, N343 is distal to the antibody binding site on spike (Fig. 4C), suggesting that N343 glycan is not involved in direct antibody binding and likely exerts changes in neutralization through effects on RBD conformational dynamics.

**FIG 4.**
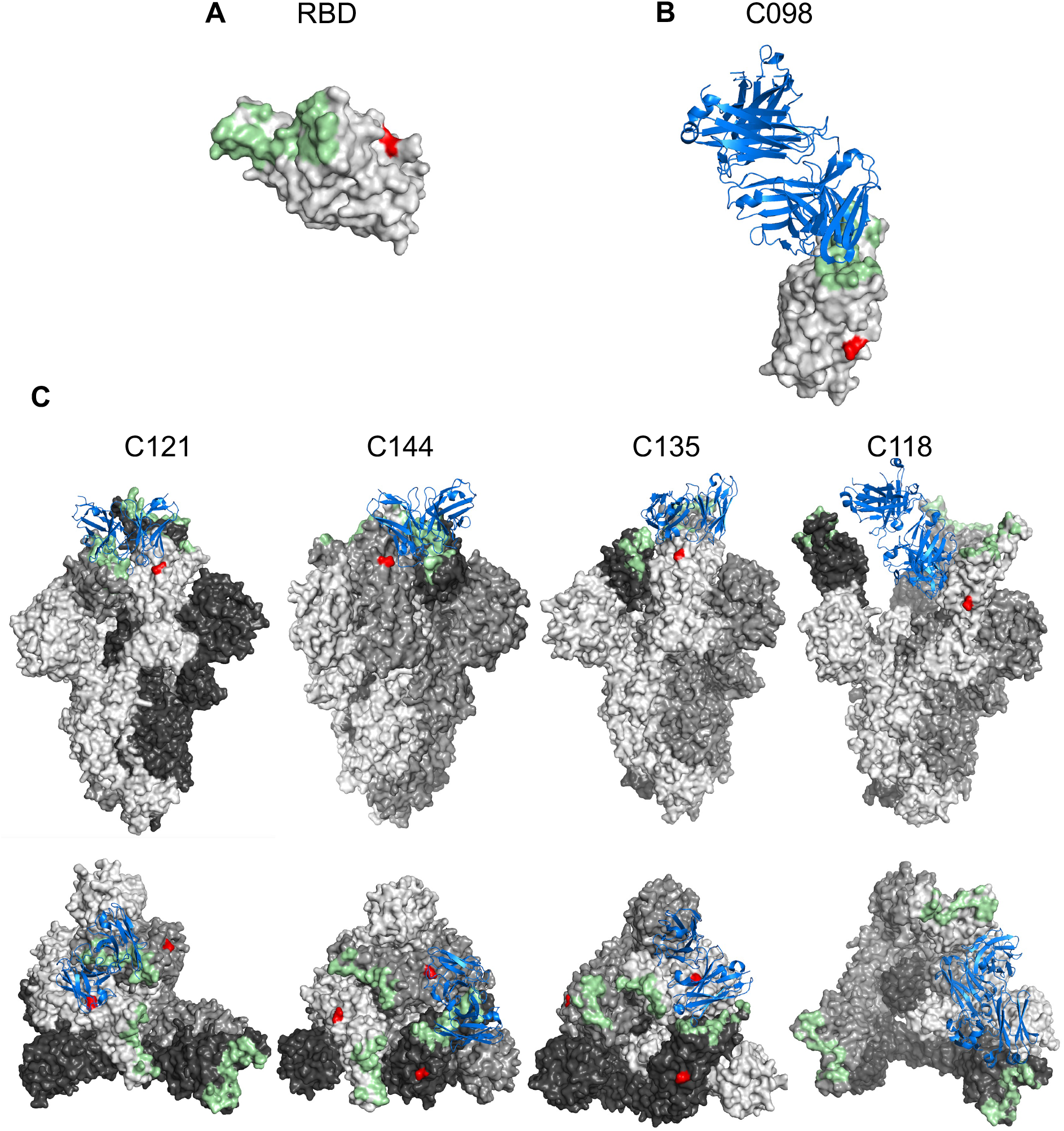
Proximity of N343 to antibody binding sites. (A) Surface representation of the receptor-binding domain (RBD) in an X-ray crystal structure (PDB ID: 7K8M). ACE2-binding site and the N343 glycosylation site are highlighted in palegreen and red, respectively. (B) Views of antibody Fab variable domains (blue) binding to spike (grey), in which each trimer subunit, is shown with distinct gray shade. ACE2-binding site and N343 are highlighted in pale green and red, respectively. The structures illustrated herein are C098 (PDB ID: 7N3I), C121 (PDB ID: 7K8X), C144 (PDB ID: 7K90), C135 (PDB ID: 7K8Z) and C118 (PDB ID: 7RKV).

Other RBD substitutions (S323A/T325A and N331D) did not alter the neutralization sensitivity to most monoclonal antibodies tested herein (Fig. S4, Fig. S5). However, the N282D substitution, which lies outside the RBD, had marginal effect on sensitivity to the class I antibodies C099, C936 and the class 3 antibody C581 (Fig. S6).

### Effect of the N343 glycan on neutralization by polyclonal SARS-CoV-2 neutralizing plasma

Given that the N343D substitution exhibited different effects on sensitivity to neutralizing monoclonal antibodies, we next asked whether this substitution affected neutralization by polyclonal antibodies present in convalescent plasma. As with the monoclonal antibodies, we compared the neutralization sensitivity of N343D pseudotyped particles with that of WT pseudotyped particles containing same amount of WT spike protein or showing the same infectivity as the N343D pseudotype, to convalescent plasma from 15 patients collected early in the COVID19 pandemic (at 1.3 months after infection) and from the same individuals ∼1 year later following subsequent vaccination (34) (36). The N343D mutant pseudotypes were more sensitive than the WT pseudotypes to convalescent plasma collected at 1.3 months after infection (Fig. 5A and Fig. S7). Indeed, the 50% neutralization titers (NT50) were a mean of 5.1-fold greater for the N343D mutant as compared to WT pseudotypes (p=0.0022) (Fig. 5B). Notably, the difference in neutralization sensitivity between N343D mutant (mean NT50=26179) and WT S pseudotypes (mean NT50=20853) was negligible (p=0.2805) when plasmas collected from the same individuals 1 year later and after subsequent vaccination were tested. Of note, these subsequently collected plasma exhibited higher neutralization potency than those collected shortly after infection (Fig. 5). We conclude that the glycan at N343 confers protection against neutralization by antibodies generated shortly after SARS-CoV-2 infection but that this effect is lost against antibodies from the same individuals who are subsequently vaccinated.

**FIG 5.**
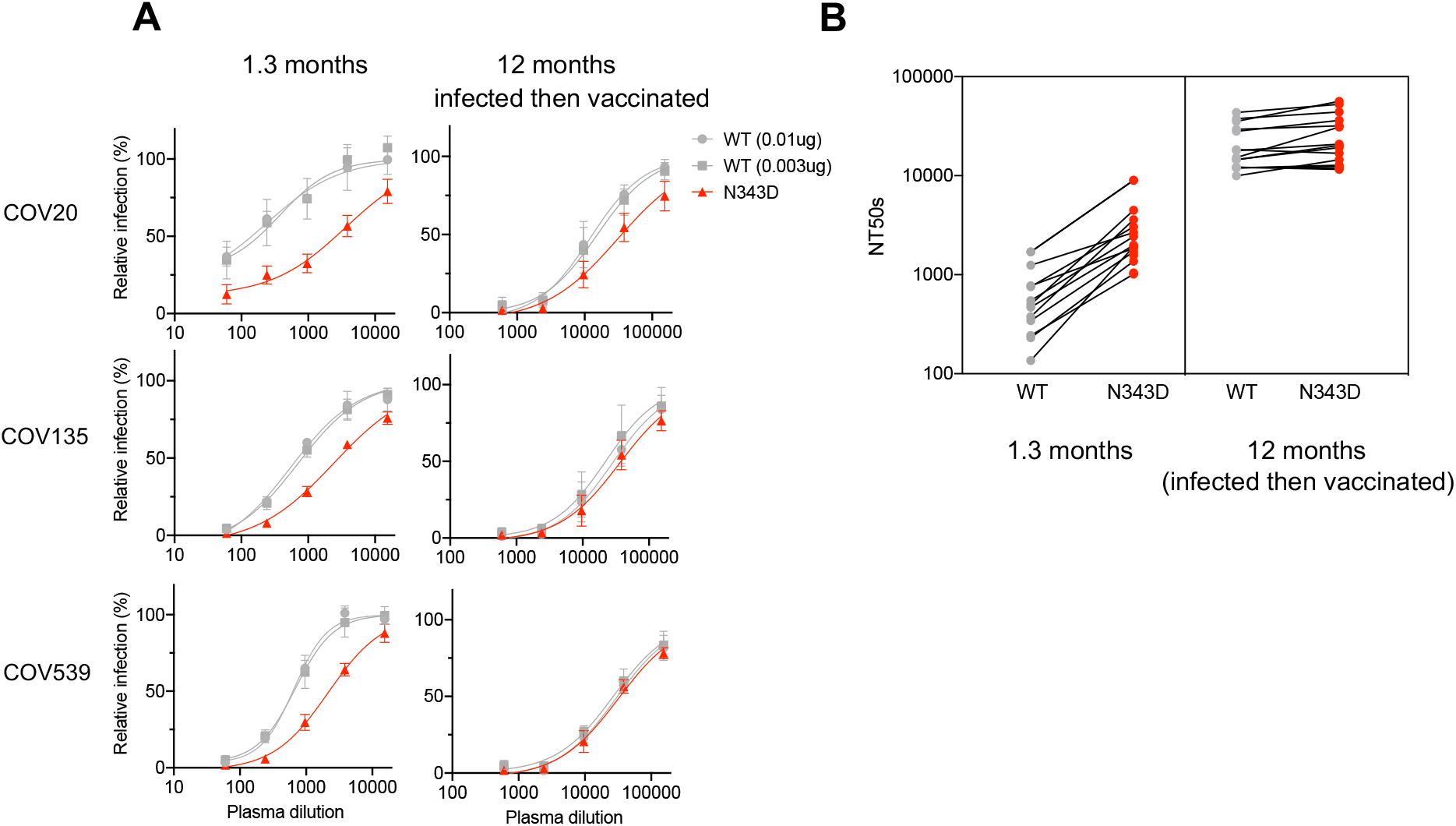
Neutralization sensitivity of N343D mutant to convalescent plasma. (A) Plasma neutralization of N343D or glycosylation site intact spike (in the furin uncleavable (R683G) background) pseudotyped virus using 293T/ACE2.cl22 target cells. The convalescent plasma samples were collected at 1.3 months and 12 months (infected then vaccinated) and diluted four-fold serially followed by incubation with viruses. As controls, glycosylation intact spike expression plasmid (WT in the R683G background) was transfected at two doses, 10 ng or 3 ng, and the neutralization sensitivity of the resulting viruses were assessed in parallel. The mean and range of two technical replicates are shown. (B) Comparison of NT_50_ values for each of the 15 convalescent plasma samples collected at 1.3 months and at 12 months (infected then vaccinated) for the N343D or glycosylation intact spike pseudotypes.

## Discussion

While SARS-CoV-2 spike pseudotyped viruses have been widely used as a surrogate to study S-mediated viral entry (28) little is known about the varying effect of S level on virions on particle infectivity and neutralization sensitivity. Previously it was reported that approximately 8 HIV-1 trimer-receptor interactions are required for HIV-1 to infect a target cell (38), a result that is broadly consistent with our finding that a small number (minimally an average of 1-2 spike per virion) is sufficient for detection of SARS-CoV-2 pseudotype infection. To achieve neutralization, monoclonal antibodies must encounter prefusion spikes and reduce the number of functional spike trimers below the threshold required for infection. We found that an increased level of S on pseudotyped virions is associated with increased infectivity but had little effect on neutralization sensitivity to monoclonal antibodies. This result suggests that monoclonal antibodies are in excess over functional spikes in a typical neutralization assay, and their measured potency is limited by their affinity, rather than by the amount of functional spike protein in a pseudotype neutralization assay.

In this study, we found that glycosylation at several sites in the spike NTD and RBD, including N61, N122, N165, N331, N343 and S323 T325 are required for full pseudotyped virus infectivity. Substitution of these glycosylation sites resulted in a reduction in pseudotype infectivity of about 10-fold or more and correlated with a reduction in spike incorporation into virions. While it is unclear precisely how glycosylation on these sites drives S incorporation into virions, glycans could in principle affect protein stability and trafficking through the secretory pathway (39). We note, however, that the glycosylation site mutations did not affect steady state levels of S in transfected cells. Compared with other glycosylation sites in the S1 domain, these sites are highly conserved among sarbecovirus S proteins (40). Likewise, the conservation of these glycosylation sites is observed among the major SARS-CoV-2 variant lineages, including Alpha, Beta, Gamma, Delta and Omicron (41). Some studies, largely employing molecular dynamics simulations, have suggested glycans on spike proteins affect the conformational dynamics of the spike’s RBD “up” and “down” states (10) (42). In particular, the glycan at N343 stabilizes RBD states in a process termed "glycan gating” (22, 43, 44). Our findings show that the N-glycan at N343 affected incorporation into virions, suggesting that the conformational state, might affect trimer assembly or stability, transit through the secretory pathway or incorporation into pseudotyped virions. The N343 glycan also affected neutralization by RBD-targeted monoclonal antibodies, and the effect was largely dependent on the class to which antibodies belong. The N343 glycan is positioned on the RBD distal to the ACE2 and class 1 antibody (C098) binding site, but increased sensitivity to class 1 antibodies (C098, C099, C936), which bind only to “up” RBDs (24), suggesting that this effect is mediated through the impact of the glycan on RBD conformation. Conversely, the N343 glycan reduced sensitivity to class 2 antibodies, including C121 and C144, both of which can bind both ‘up’ and ‘down’ RBDs. Cryo-EM structures (24) (Fig. 4) show that N343 is located proximally to the C121 and C144 antibody bound to the neighboring subunit, in a manner that might interfere with antibody binding. Overall, our results are consistent with a model in which N343 glycan affects sensitivity to class 1 and class 2 antibodies by affecting the RBD conformational dynamics (43) and also potentially by sterically hindering class 2 antibody binding into neighboring RBDs in the “down” conformation (Fig. 4).

The N343 glycan is proximal to the binding sites of class 3 antibodies, and the N343D substitution had a range of effects on sensitivity to class 3 antibodies. Three clonally related antibodies, C032, C080 and C952, were unaffected by the N343D substitution, while substantial but opposing effects were seen for two others, C135 and C581. In the case of C135, removal of the glycan reduced the fraction of virions that resisted neutralization at high antibody concentration. A possible explanation for this phenomenon is that the N343 site is heterogeneously glycosylated, and some subfraction of the glycans occlude the C135 binding site. For C581, removal of the N343 glycan had the opposite effect, reducing neutralization sensitivity. In this case it seems likely C581 mimics the properties of two previously described cross-reactive class 3 antibodies, S309 and SW186, for which the N343 glycan constitutes part of the antibody binding site (25) (26). Finally, for the two class 4 antibodies, the glycan at N343 affected the character of the neutralization curve. Since in these cases the antibody binding site is on the opposite face of the RBD to that of the glycan, it is likely that these effects are mediated through alteration of spike conformational dynamics and exposure of the class 4 epitope, which is shielded in the RBD ‘down’ conformation.

The N343 glycan reduced neutralization by convalescent plasma collected at 1.3 months after infection, echoing the effects on sensitivity to the C121, C144 (class 2) and C135 (class 3) antibodies. Conversely, neutralization by convalescent plasma from the same individuals (collected at 12 months following subsequent vaccination) was unaffected by the N343 glycan. That neutralizing antibodies are relatively sensitive to glycan-mediated protection early after infection may be a reflection of the fact that the initial neutralizing response is based to a large extent on class 2 antibodies that are easily escaped (29, 34). Subsequent increases in antibody affinity and neutralizing potency and a shift in the RBD epitopes that are recognized, following months of affinity maturation and boosting by vaccination (7, 35, 36, 45) results in neutralizing antibodies that are mostly unaffected by the N343 glycan. In sum, while the SARS-CoV-2 N343 glycan affects both spike conformation and neutralization sensitivity shortly after infection, antibody evolution confers sufficient potency and breadth to combat glycan-mediated immune evasion.

## Acknowledgements

We thank members of the Bieniasz and Hatziioannou laboratories for helpful discussions. This work is supported by grants from NIAID: P01 AI165075 (P.D.B. and T.H.), R37AI64003 (P.D.B.), R01AI157809 (P.D.B.) and R01AI78788 (T.H.). P.D.B. and M.C.N. are Howard Hughes Medical Institute (HHMI) Investigators. The funders had no role in study design, data collection and analysis, decision to publish or preparation of the manuscript. This Article is subject to HHMI’s Open Access to Publications policy. HHMI laboratory heads have previously granted a non-exclusive CC BY 4.0 license to the public and a sublicensable license to HHMI in their research articles. Pursuant to those licenses, the author-accepted manuscript of this article can be made freely available under a CC BY 4.0 license immediately on publication.

## Author Contributions

F.Z., T.H. and P.D.B. conceived the study. F.Z. constructed spike mutants, expressed proteins, performed Western Blot analysis and the pseudovirus neutralization assays. F.S. provided the wild-type spike plasmid as well as technical support. F.M., C.G., M.C. and M.C.N provided monoclonal antibodies. C.G., M.C. and M.C.N. provided convalescent plasma from individuals with COVID-19. P.D.B., T.H. and M.C.N. supervised the work. F.Z., T.H. and P.D.B. wrote the manuscript with input from all co-authors.

## Materials and Methods

### Antibodies and recombinant HIV p24

Antibodies used here are anti-SARS CoV-2 spike antibody [1A9] (Genetax, GTX632604) and anti-HIV capsid p24 (183-H12-5C, NIH AIDS Research and Reference Reagent Program). Secondary antibodies included goat anti-mouse conjugated to IRDye 800CW or IRDye 680 for Western blot analysis. The recombinant HIV p24 protein was purchased from abcam (ab43037).

### Plasmid construction

The plasmid expressing C-terminally truncated, human-codon-optimized SARS-CoV-2 spike protein (pSARS-CoV-2_Δ19_) has been previously described (28). Using this plasmid as the template, asparagine at N-linked glycosylation sites was replaced by aspartate by overlap-extension PCR amplification with primers that incorporated the corresponding nucleotide substitutions. O-linked glycosylation sites in the RBD region (S323, T325) were replaced by alanine using the same strategy. The purified PCR products were then inserted into the pCR3.1 expression vector with NEBuilder® HiFi DNA Assembly. Some mutations within or adjacent to the RBD region, including N282D, N331D, N343D, and S323A T325A, were also introduced in spike bearing the R683G substation which impairs the furin cleavage site and enhances particle infectivity.

### Cell lines

Human embryonic kidney HEK-293T cells (ATCC CRL-3216) and the derivative expressing ACE2, ie 293T/ACE2.cl22, were maintained in Dulbecco’s Modified Eagle Medium (DMEM) supplemented with 10% fetal bovine serum (Sigma F8067) and gentamycin (Gibco). All cell lines used in this study were monitored periodically to ensure the absence of retroviral contamination and mycoplasma.

### Generation of clarified SARS-CoV-2 pseudotyped virions and infectivity measurement

To generate HIV-1/NanoLuc-SARS-CoV-2 pseudotyped virions, two million 293T cells in 10-cm dish were transfected with 7.5 μg of HIV-1 proviral plasmids expressing NanoLuc along with increasing amounts (0 ng, 2 ng, 3.9 ng, 7.8 ng, 15.6 ng, 31.2 ng, 62.5 ng, 125 ng, 250 ng, 500 ng, or 1000 ng) of WT or 1000 ng of glycosylation site mutant SARS-CoV-2 expression plasmids (pSARS-CoV-2_Δ19_). In transfection experiment to generate viruses bearing R683G substitution, 1 ng, 3 ng, 10 ng, 30 ng, 100 ng, and 300 ng of expression plasmid were used instead. The total amount of DNA was held constant by supplementing the transfection with empty expression vector. Cells were harvested at 48 hr post transfection and subjected to Western blot analysis. Virus-containing supernatant was filtered (0.2 μm), and, to remove cell debris, clarified by Lenti-X (TaKaRa). Particle infectivity was measured as previously described (28). Briefly, viral stocks were three-fold serially diluted and added to 293T/ACE2 cl.22 in 96-well plates. Cells were then harvested at 48 hr post infection for measuring NanoLuc activity (Promega). The number of spike trimers per virion was estimated using the following formula: S ng/ml/78.3 x 1500/(p24 ng/ml/24)/3.

### Western blot analysis

Cell lysates were separated on NuPage Novex 4-12% Bis-Tris Mini Gels (Invitrogen). Proteins were blotted onto nitrocellulose membranes. Thereafter, the blots were probed with primary antibodies and followed by secondary antibodies conjugated to IRDye 800CW or IRDye 680. Fluorescent signals were detected and quantitated using an Odyssey scanner (LI-COR Biosciences).

### S-6P-NanoLuc protein purification

To express S-6P-NanoLuc proteins, Expi293 cells were transfected with S-6P-NanoLuc expression plasmids using ExpiFectamine 293 (ThermoFisher Scientific). Four days later, the supernatant was harvested and loaded on Ni-NTA agarose and, after thorough wash, S-6P-NanoLuc proteins were released after HRV 3C protease treatment.

### Neutralization assays

To measure neutralizing activity of monoclonal antibodies, serial dilutions of antibodies beginning at 3 μg/ml were four-fold serially diluted in 96-well plates over seven dilutions. To determine the neutralizing activity in convalescent plasma, the initial dilution started at a 1:30 (for plasma at 1.3 months) or a 1:150 (for plasma at 12 months). Thereafter, SARS-CoV-2 spike pseudotyped viruses were incubated with monoclonal antibodies or the convalescent plasma for 1 hr at 37°C in 96-well plates. The mixture was then added to 293T/ACE2cl.22 target cells seeded one day prior to infection so the final starting dilutions were 1.5 μg/ml for monoclonal antibodies and 1:60 or 1:300 for plasma. Cells were then harvested 48 hours post infection for NanoLuc luciferase assays.

### Human plasma samples and monoclonal antibodies

Monoclonal antibodies C022, C032, C080, C098, C099, C118, C121, C135, C144, C581, C936, and C952 used in this study were previously reported (24, 34-37). The human convalescent plasma samples (COVs) were obtained under protocols approved by Institutional Review Boards at Rockefeller University.

## Figures

**FIG S1.**
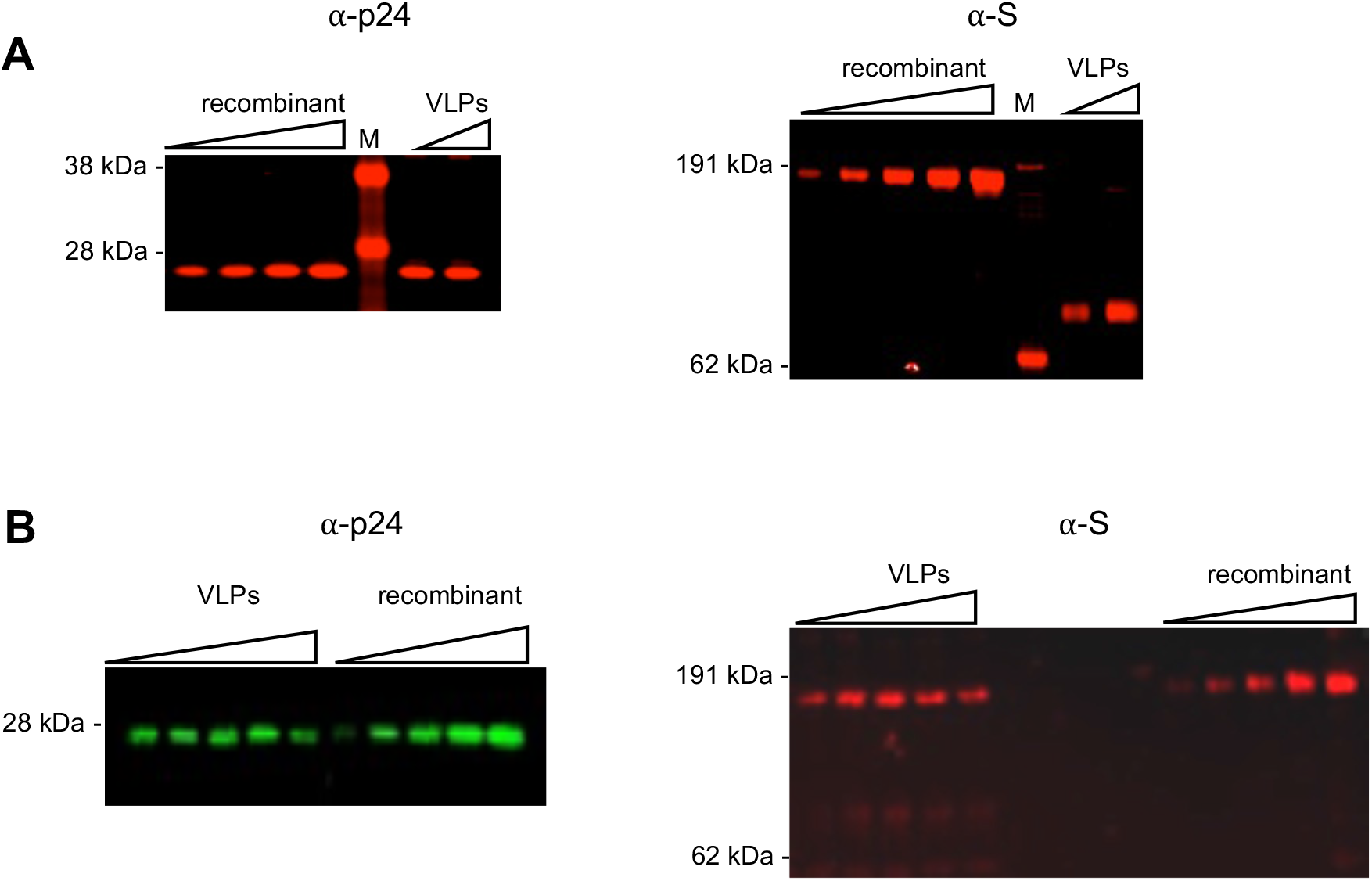
Western blot analysis of virions, using recombinant proteins as standard. (A) Western blot analysis of virions pelleted through 20% sucrose from 100 μl supernatant harvested at 48 hours after transfection with 0.0625 μg or 0.5 μg of wild-type spike expression plasmid along with envelope-deficient HIV-1 proviral plasmid expressing NanoLuc luciferase. The blot was probed with an anti-p24 antibody and recombinant HIV p24 protein was used as a standard (1.0 ng, 2.0 ng, 4.0 ng, or 8.0 ng per lane) on the left, or with anti-Spike antibody using recombinant S-6P-nanoLuc as a standard (0.25 ng, 0.5 ng, 1.0 ng, 2.0 ng, or 4.0 ng per lane) on the right. Representative of two independent experiments. (B) Western blot analysis of virions pelleted through 20% sucrose from 100 μl supernatant harvested at 48 hours after transfection with 0.008 μg, 0.024 μg, 0.073 μg, 0.22 μg, or 0.67 μg of wild-type spike expression (furin uncleavable R683G background) along with envelope-deficient HIV-1 proviral plasmid expressing NanoLuc luciferase. The blot was probed with anti-p24 antibody using recombinant HIV p24 protein as standard (0.5 ng, 1.0 ng, 2.0 ng, 4.0 ng, or 8.0 ng per lane) on the left, or with anti-Spike antibody using recombinant S-6P-nanoLuc as standard (0.125 ng, 0.25 ng, 0.5 ng, 1.0 ng, or 2.0 ng per lane) on the right. Representative of two independent experiments.

**FIG S2.**
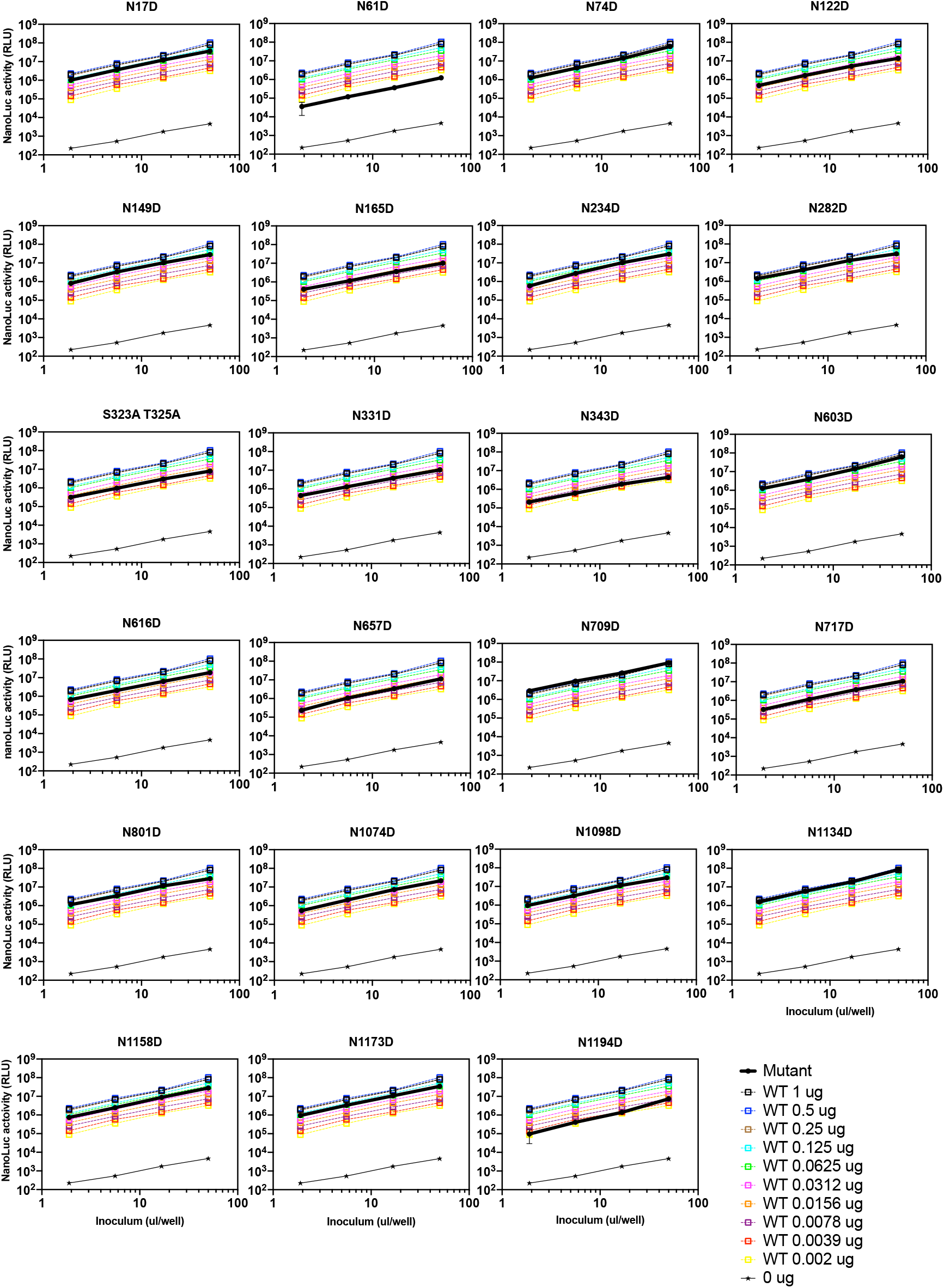
The impact of glycosylation site mutations on particle infectivity. Infectious virion measurements of for pseudotypes bearing glycosylation site mutant spike proteins (bold black lines, 1 μg transfected S expression plasmid), compared with as well as WT S pseudoytpes (dashed lines) collected from 293T cells transfected with various amounts of spike expression plasmids. Infection was quantified by measuring NanoLuc luciferase activity (RLU). Virus generated in the absence of S (0 μg), shown in thin black line, was used as a background control. 293T/ACE2.cl22, as target cells, were infected with the indicated volumes of pseudotyped viruses in 96-well plates and harvested 48 hours post infection for NanoLuc luciferase assay. The mean and range deviation from two technical replicates are shown.

**FIG S3.**
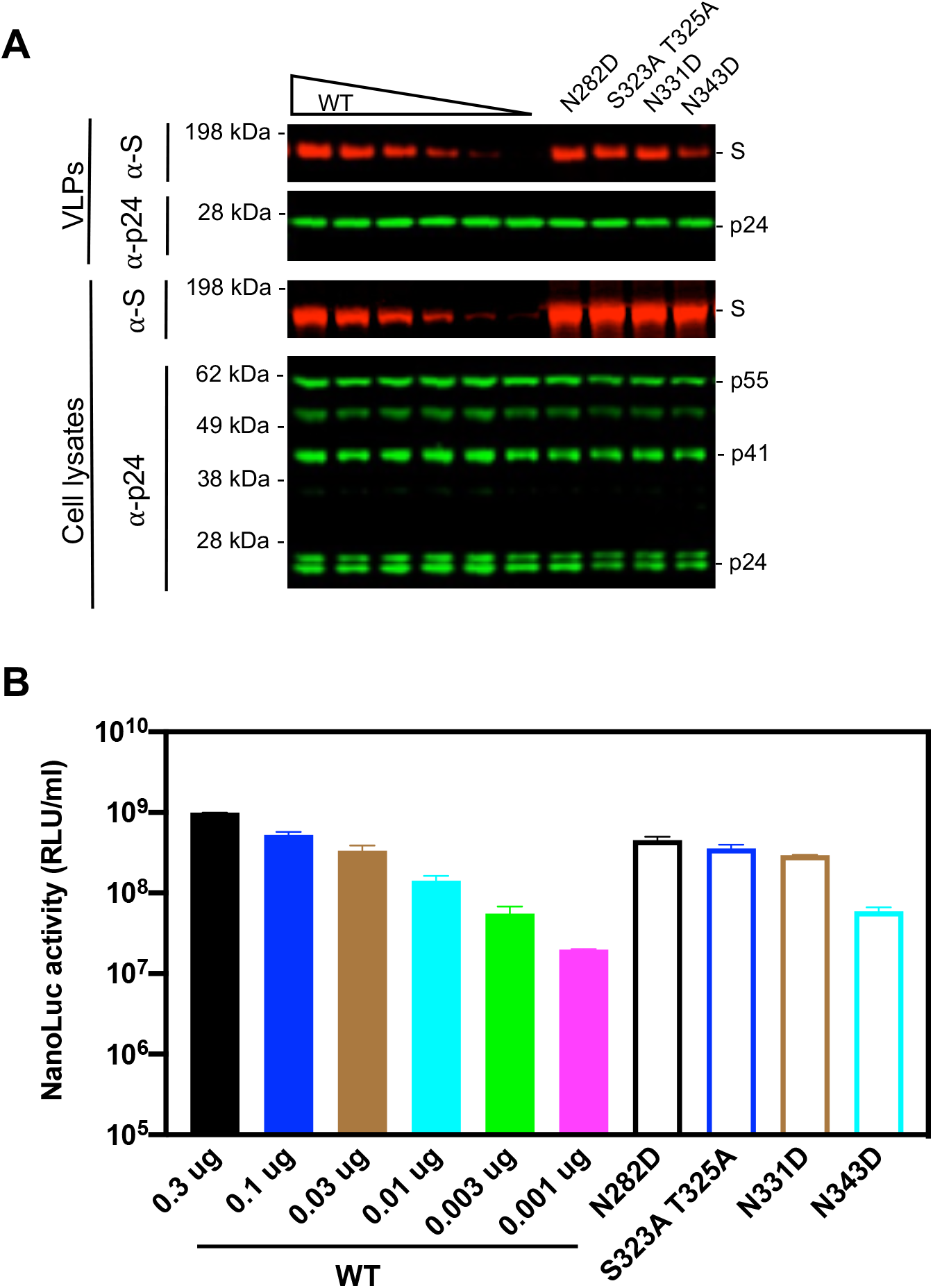
The impact of glycosylation site mutations in the furin uncleavable (R683G) background on spike incorporation and particle infectivity. (A) Western blot analysis of 293T cell lysates (lower panel) or virions (upper panel) at 48 hours after transfection with various amounts of glycosylation site intact spike expression plasmid (R683G background), or 1 μg of glycosylation site mutants (N282D, S323A T325A, N331D, or N343D) along with envelope-deficient HIV-1 proviral plasmid expressing NanoLuc. (B) Infectivity was quantified by measuring NanoLuc luciferase activity (RLUs) following infection of 293T expressing ACE2 (293T/ACE2.cl22) in 96-well plates with pseudotyped viruses as depicted in (A). The mean and range of two technical replicates are plotted.

**FIG S4.**
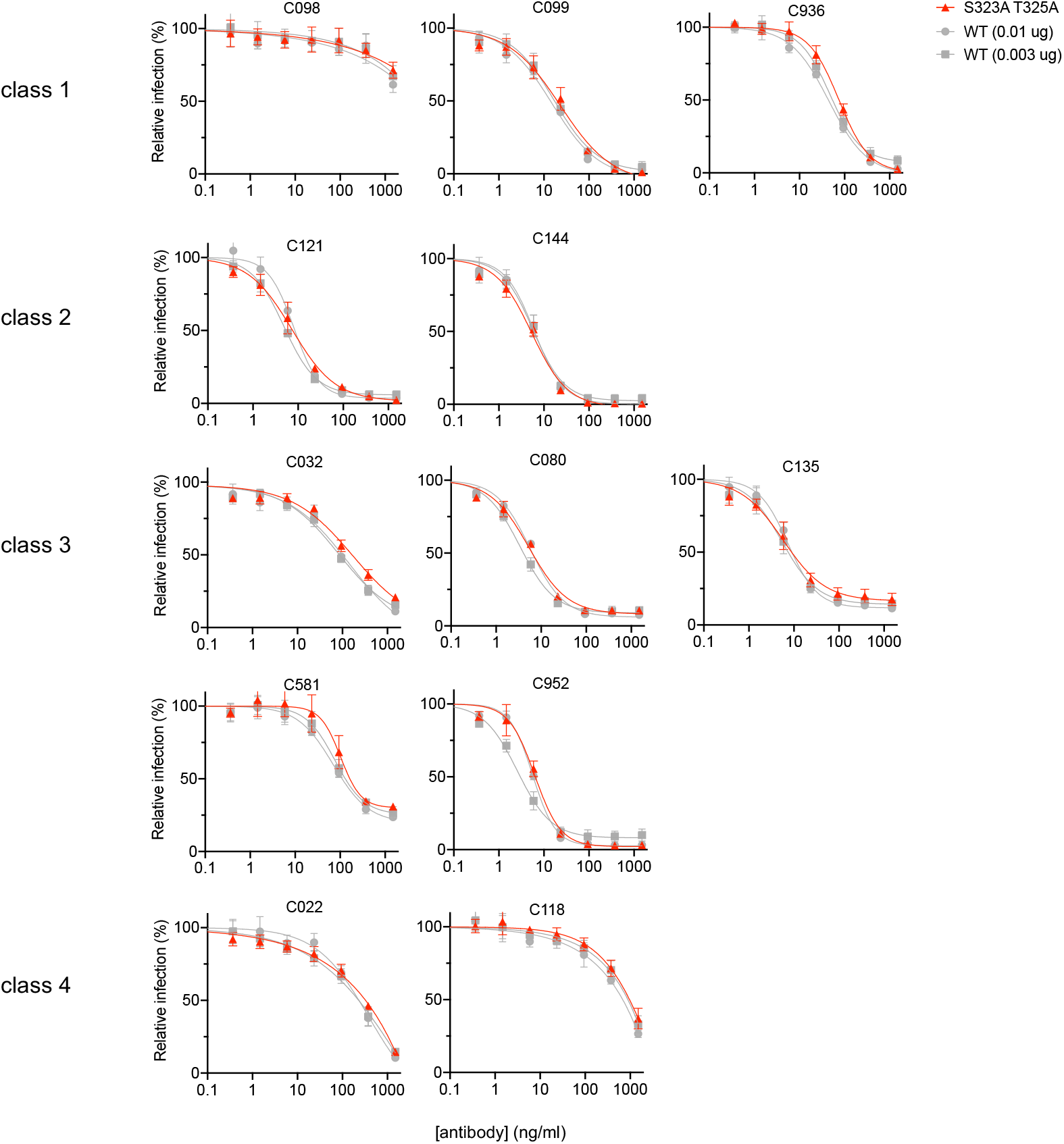
Mutations of the O-linked glycosylation sites at 323 and 325 in the RBD (S323A T325A) have marginal effect on neutralization sensitivity. Neutralization of glycosylation site mutant S323A T325A pseudotyped virus infection in the presence of the indicated concentrations of a panel of monoclonal antibodies, including class 1 (C098, C099, and C936), class 2 (C121 and C144), class 3 (C032, C080, C135, C581, and C952), and class 4 (C022 and C118) antibodies. As controls, glycosylation intact spike expression plasmid (WT in the furin uncleavable R683G background) was transfected at two doses, 10 ng or 3 ng, and the resulting viruses were assessed for neutralization sensitivity in parallel. The mean and range of two technical replicates are shown.

**FIG S5.**
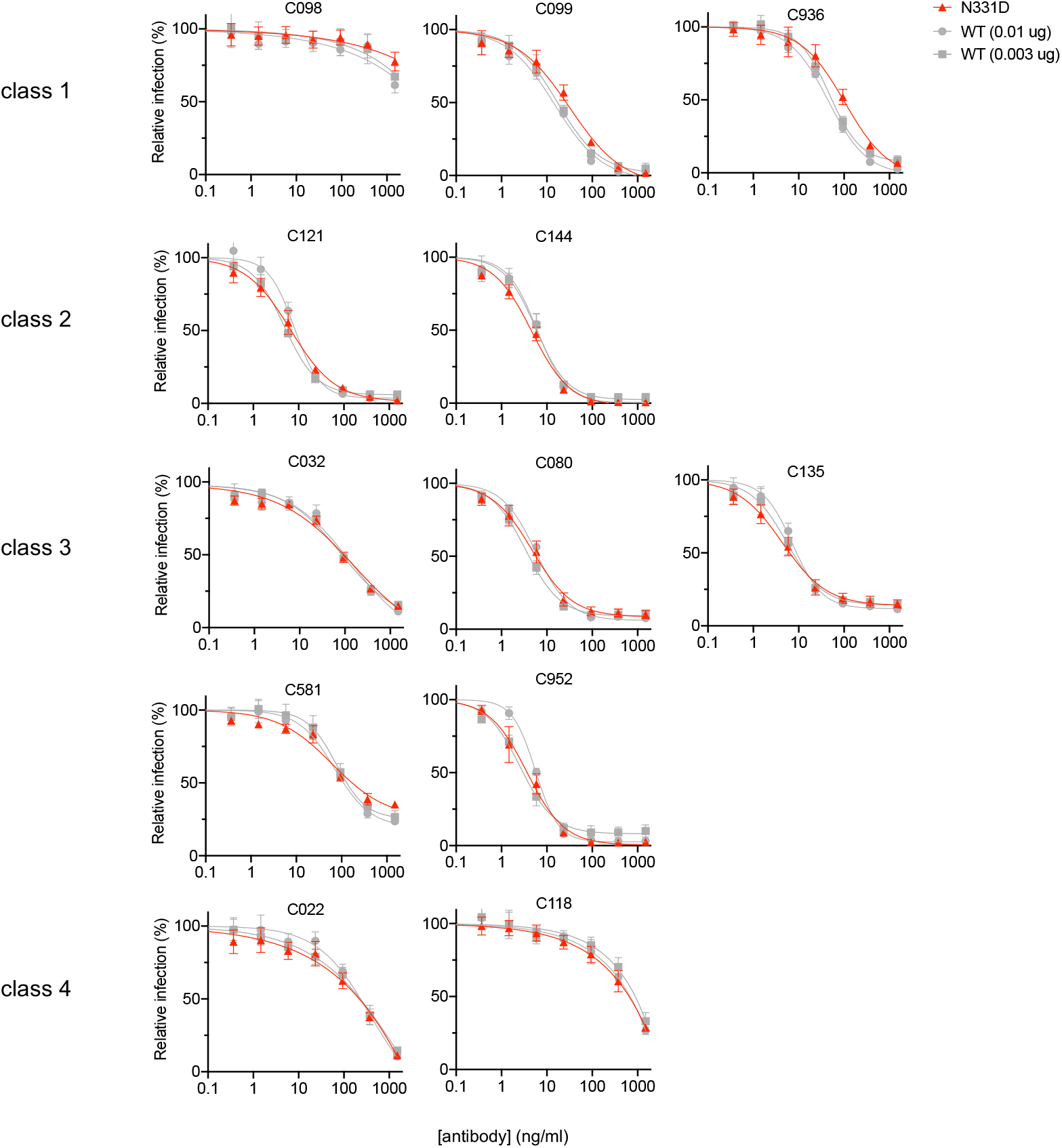
Effect of glycosylation at N331 on neutralization sensitivity. Neutralization of glycosylation site mutant N331D pseudotyped virus infection in the presence of the indicated concentrations of a panel of monoclonal antibodies, including class 1 (C098, C099, and C936), class 2 (C121 and C144), class 3 (C032, C080, C135, C581, and C952), and class 4 (C022 and C118) antibodies. As controls, glycosylation intact spike (WT in the furin uncleavable R683G background) was transfected at two doses, 10 ng or 3 ng, and the resulting viruses were assessed for neutralization sensitivity in parallel. The mean and range of two technical replicates are shown.

**FIG S6.**
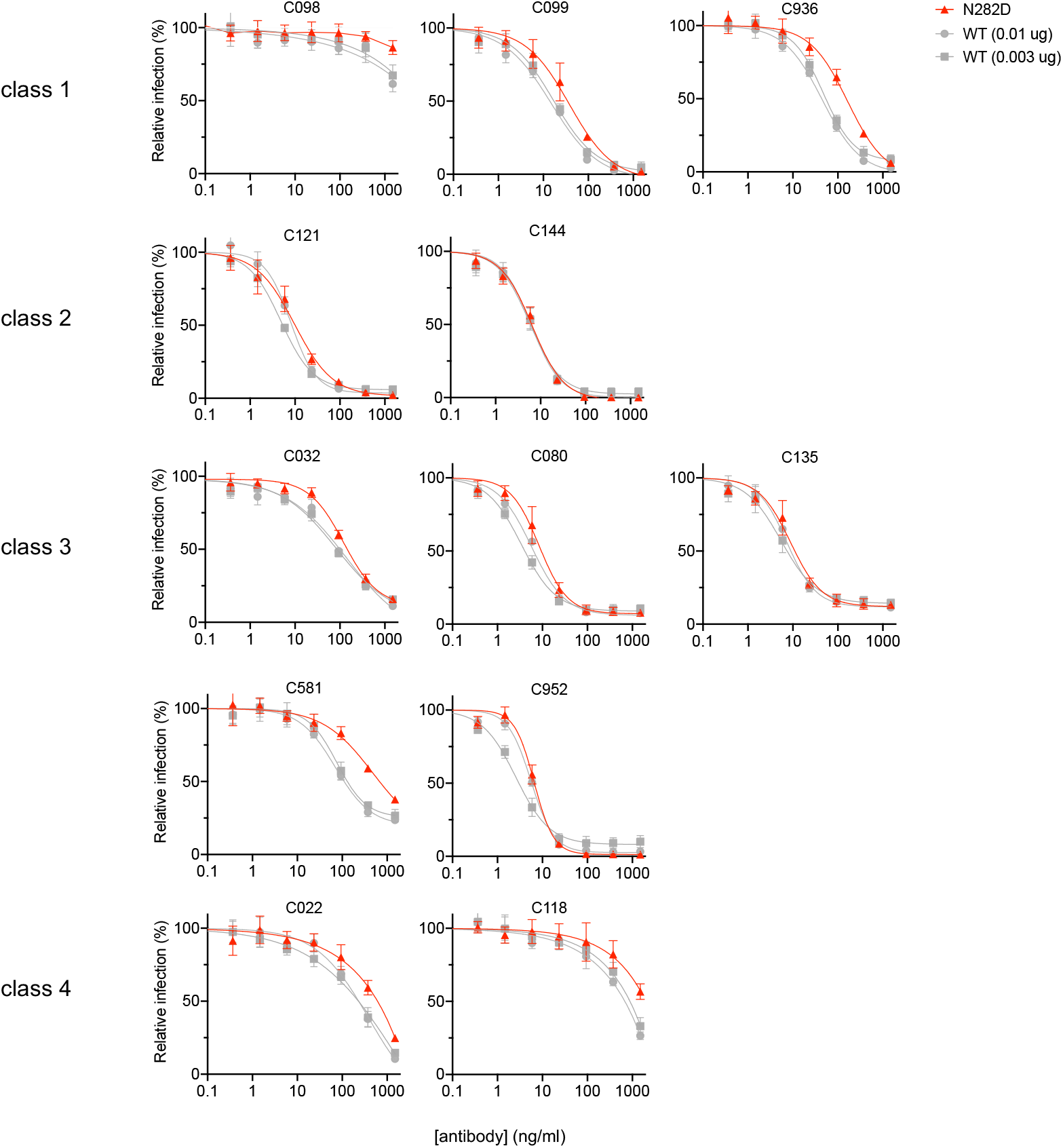
Effect of glycosylation at N282 on neutralization sensitivity. Neutralization of glycosylation site mutant N282D pseudotyped virus infection in the presence of the indicated concentrations of a panel of monoclonal antibodies, including class 1 (C098, C099, and C936), class 2 (C121 and C144), class 3 (C032, C080, C135, C581, and C952), and class 4 (C022 and C118). As controls, glycosylation intact spike (WT in the furin uncleavable R683G background) was transfected at two doses, 10 ng or 3 ng, and the resulting viruses were assessed for neutralization sensitivity in parallel. The mean and range of two technical replicates are shown.

**FIG S7.**
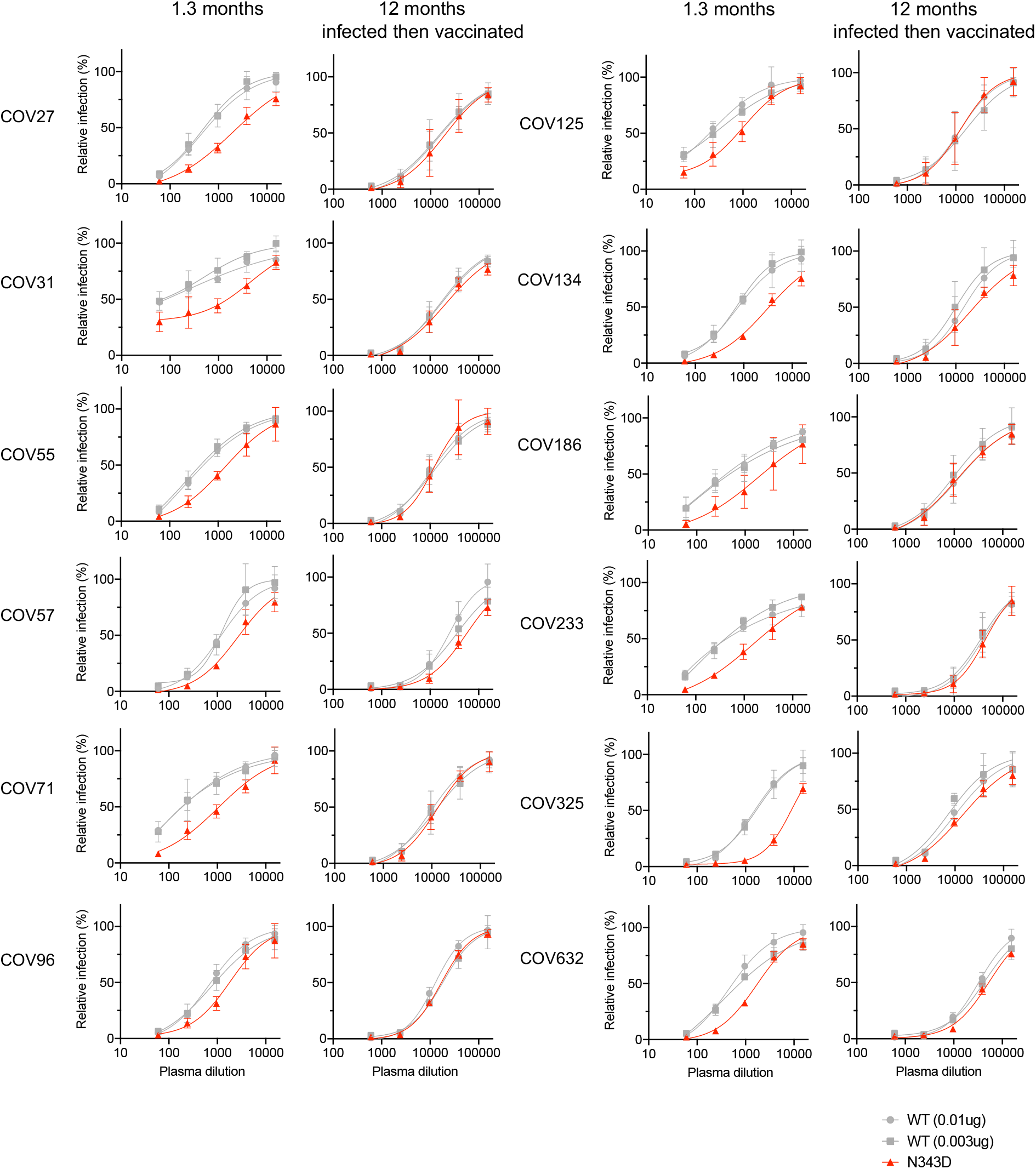
Neutralization sensitivity of N343D mutant to convalescent plasma. Additional examples of plasma neutralization of N343D or glycosylation site intact spike (in the furin uncleavable R683G background, same as FIG 5) pseudotyped virus using 293T/ACE2.cl22 target cells. The mean and range of two technical replicates are shown.

